# Sleep consolidation potentiates skill maintenance

**DOI:** 10.1101/2023.10.26.564031

**Authors:** Agustin Solano, Gonzalo Lerner, Guillermina Griffa, Alvaro Deleglise, Pedro Caffaro, Luis Riquelme, Daniel Perez-Chada, Valeria Della-Maggiore

## Abstract

Contrary to its well-established role in declarative learning the impact of sleep on motor memory consolidation remains a subject of debate. Motor learning involves skill acquisition and skill maintenance, two critical aspects in achieving movement precision. Current literature suggests that whereas motor skill acquisition benefits from sleep, consolidation of skill maintenance depends solely on the passage of time. This has led to the proposal that skill maintenance may be an exception to other types of memories. Here, we address this ongoing controversy in humans through three comprehensive experiments. We found that when training occurs throughout the day consolidation of skill maintenance proceeds independently of sleep. However, when it takes place closely before bedtime so that sleep aligns with the memory stabilization window, a 30% memory enhancement emerges along with a distinct modulation of neural markers of sleep consolidation. Our findings reconcile seemingly conflicting perspectives on the active role of sleep in procedural motor learning, and offer the potential to accelerate motor recovery in rehabilitation programs through the synchronization of training sessions with sleep.

## Introduction

The role of sleep in declarative memory consolidation is well established. Sleep consistently enhances memory retention in a variety of declarative learning paradigms including but not limited to face recognition (e.g., Wagner et al., 2007), free recall (e.g., Lahl et al., 2008; Mednick et al., 2008), and paired associates (e.g., Tucker et al., 2006; Talamini et al., 2008; Tucker & Fishbein, 2008; Lau et al., 2010; Payne et al, 2012; Diekelmann et al., 2012). In contrast, the contribution of sleep to non-declarative motor learning is more equivocal and, at first sight, appears to vary remarkably with the experimental paradigm. Motor learning encompasses skill acquisition — the incorporation of new motor programs for precise movement execution-, and skill maintenance —the ability to recalibrate pre-existing motor programs under changing environmental or internal conditions (Krakauer et al., 2019). These aspects of learning often referred to as motor skill learning (MSL) and sensorimotor adaptation (SMA) respectively, are interdependent and support a wide array of human actions such as serving a tennis ball and making the necessary adjustments to maintain precision when playing in gusty conditions.

One of the most popular experimental paradigms to study MSL in the laboratory involves performing a sequence of finger movements on a keyboard using the non-dominant hand. SMA has also been studied extensively using experimental paradigms in which subjects learn to reach targets while facing visual perturbations (such as an optical rotation) or proprioceptive perturbations (like force fields), which alter sensorimotor coordination and, hence, the trajectory of the arm. Substantial evidence underscores the significance of non-rapid eye movement (NREM) sleep in MSL memory stabilization (Rickard et al., 2008; Brawn et al., 2010; Nettersheim et al., 2015) or in the emergence of overnight offline gains when the sequence is encoded explicitly (Robertson et al., 2004; Nishida & Walker, 2007; Doyon et al., 2009; Diekelmann & Born, 2010; Albouy et al., 2013; Breton & Robertson, 2017). In contrast, the available evidence on SMA suggests that the consolidation of this type of motor memory is independent of sleep. Specifically, Donchin et al. (2002) have demonstrated that one night of sleep deprivation after learning an SMA task that requires reaching to targets through a force field does not impair memory retention. Thürer et al. (2018) obtained comparable results using the same experimental paradigm with a period of wakefulness or sleep that followed learning. In the same vein, Doyon et al. (2009) and Debas et al. (2010) have shown similar levels of memory retention when visuomotor adaptation is followed by an equivalent period of sleep or wakefulness. Based on these findings, it has been argued that, unlike MSL, the consolidation of SMA relies exclusively on the passage of time (Brodt et al., 2023).

At first sight, this discrepancy between declarative learning and MSL on one side, and SMA on the other side suggests the presence of different mechanisms supporting memory consolidation depending on the memory system and/or the experimental paradigm. It is noteworthy, however, that the majority of the studies reviewed above focused on tracking memory retention after a time interval that includes –or not-a period of sleep, but tended to overlook *the temporal gap between training and bedtime* as a relevant factor. The close proximity between these events is indeed a strong modulator of declarative and motor sequence memories (Barrett & Ekstrand, 1972; Benson & Feinberg, 1977; Gais et al., 2006; Talamini et al., 2008; Payne et al., 2012; Doyon et al., 2009; Van Der Werf et al., 2009; Holz et al., 2012; Truong et al., 2023). Thus, it is possible that like MSL and declarative tasks, SMA also benefits from sleep when the latter occurs closely after training. We speculate that this phenomenon may result from the temporal overlap between memory stabilization initiated during learning and the neurophysiological processes ongoing during NREM sleep.

In this study, we propose that like motor skill acquisition, motor skill maintenance consolidates both during wakefulness and sleep. To test this hypothesis, we conducted a series of experiments (Figure 1) using a well-established SMA paradigm involving visuomotor adaptation to an optical rotation (e.g. Villalta et al., 2015; Lerner et al., 2020; Albert et al., 2022). In Experiment 1, we investigated the impact of sleep on memory retention when training occurred far from bedtime. Based on the revised literature (Donchin et al., 2002; Doyon et al., 2009; Debas et al., 2010; Thürer et al., 2018), we predicted that sleep would not benefit SMA memory under these conditions. Subsequently, in Experiment 2, we sought to establish the time course of memory stabilization during wakefulness using an anterograde interference protocol. This experiment aimed to determine the specific time window during which sleep might be most effective in modulating SMA memory. Building upon the insights from Experiment 2, in Experiment 3 we examined the impact of sleep on long-term memory retention when training occurred immediately before bedtime. Furthermore, to ascertain whether sleep operates through an active mechanism -as opposed to merely providing protection against interference-we used EEG to quantify neural markers of consolidation well-established in the declarative literature, namely, the density of fast sleep spindles and their coupling with slow oscillations (Maingret et al., 2016; Ladenbauer et al., 2017; Helfrich et al., 2018; Muehlroth et al., 2019; Navarro-Lobato & Genzel, 2019). We predicted that NREM sleep would benefit SMA through an active mechanism only when it overlaps with the stabilization window.

**Figure 1.**
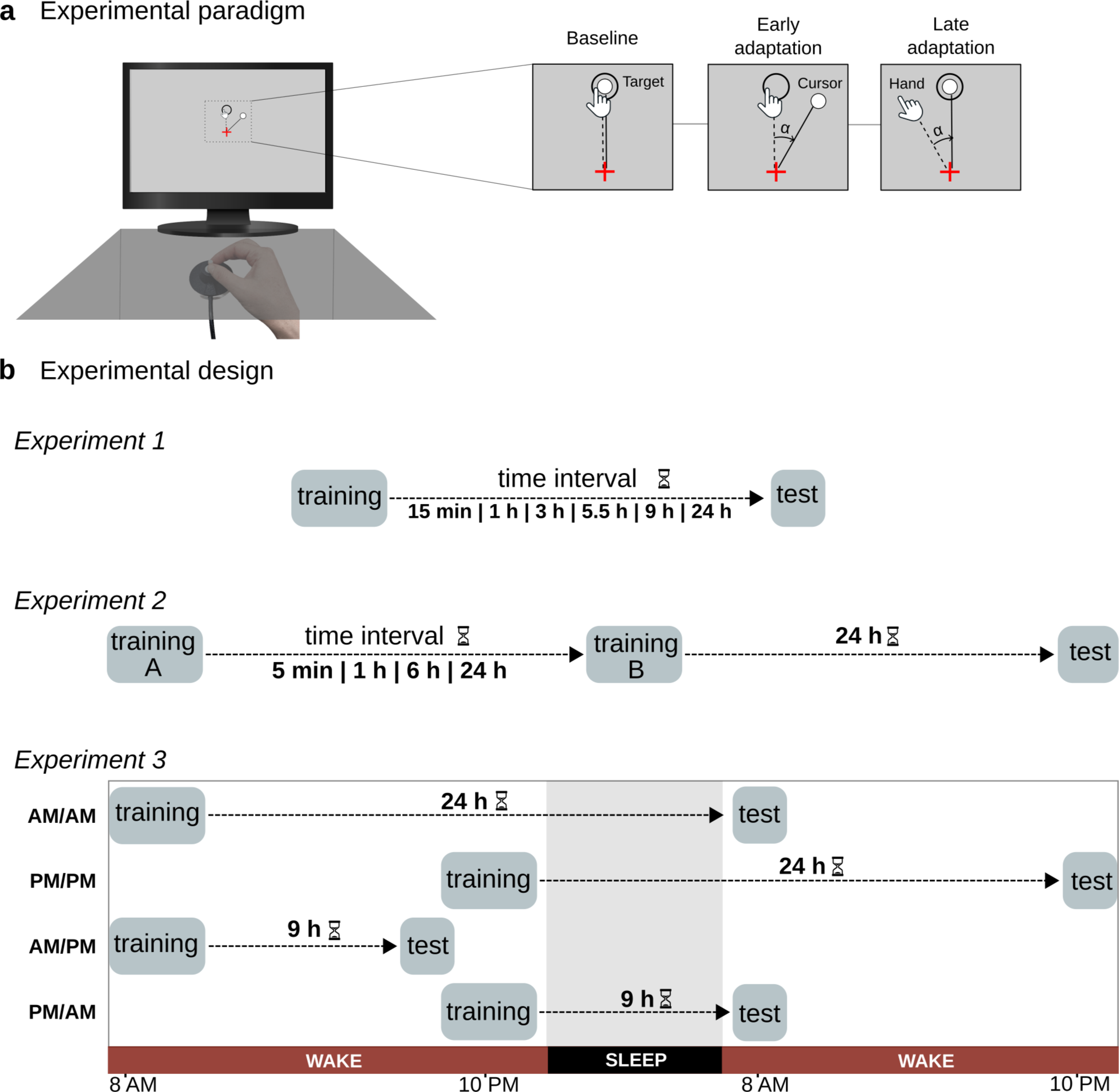
Experimental paradigm and experimental design. a) SMA experimental paradigm. Subjects sat on a chair and performed center-out movements to one of eight visual targets using a cursor controlled with a joystick operated with their right -dominant-hand. One cycle was composed of 8 trials (one per target), and 11 cycles composed a block. The vision of the hand was occluded. The inset represents the visual display of the computer screen, showing the relationship between the hand and cursor trajectory before (baseline), and during the early and late stages of adaptation to an optical rotation (α). b) Experimental design. *Experiment 1*: Effect of sleep on SMA when training occurs far from bedtime. Six different groups of participants were trained on the SMA paradigm (*training*) and their memory retention was tested after variable time intervals post-learning (*test*): 15 min, 1 h, 3 h, 5.5 h, 9 h, or 24 h. During *test*, error-clamp (EC) trials were used to assess memory retention by simulating ‘straight’ paths to targets with controlled variability. Training took place throughout the daytime. Only the 24 h group underwent a full night of sleep between training and test. *Experiment 2*: Time course of SMA memory stabilization during wakefulness. Four groups of participants adapted to two opposing optical rotations denoted as A and B (*training A* and *training B*), separated by either 5 min, 1 h, 6 h, or 24 h. An additional control group only adapted to B. Memory retention was assessed with EC trials 24 h after *training B*, during the *test* session. *Experiment 3*: Effect of sleep on SMA when training occurs immediately before bedtime. Two groups of volunteers were trained on the SMA task (*training*) and polysomnographic EEG recordings were performed overnight. The AM/AM group slept far from training (∼14 h after training on the SMA task), whereas the PM/PM group went to sleep immediately after training (∼10 min). Memory retention was assessed 24 h after training with EC trials (*test*). To rule out a possible circadian effect due to the time of *test*, one additional group (AM/PM), trained AM and was tested PM, ∼9 h later, without intermediate sleep, whereas another group (PM/AM) trained PM and was tested overnight (AM), ∼9 h later.

## Results

### Sleep does not benefit motor skill maintenance when training occurs far from bedtime

In contrast to declarative and motor sequence learning, the consolidation of SMA memory has consistently been shown to depend exclusively on the passage of time (Donchin et al., 2002; Doyon et al., 2009; Debas et al., 2010; Thürer et al., 2018). In this study, we hypothesized that the apparent lack of a sleep benefit can be attributed to the considerable temporal gap between training and bedtime. In Experiment 1 we tested this hypothesis by quantifying the evolution of memory retention through a 24 h period when the time interval between the two events was not controlled for. The obtained curve of forgetting allowed us to determine whether sleep potentiated or simply stabilized memory. Six groups of volunteers (n=20-25 per group) performed a visuomotor adaptation SMA task in which a visual perturbation in the form of an optical rotation (30-degree counterclockwise; CCW) was applied to a cursor while subjects made pointing movements to one of eight visual targets with the right –dominant-hand (Figure 1a; Villalta et al., 2015; Lerner et al., 2020). With time, participants learned to recalibrate the visuomotor mapping by moving in the opposite direction. Using a between-subject design, we tracked memory retention at 6 time intervals post-training through a 24 h window: 15 min, 1 h, 3 h, 5.5 h, 9 h, and 24 h (Figure 1b, top panel). Critical to the aim of Experiment 1, the time of training was not fixed either within group or across groups. For example, subjects in the 15 min group were trained sometime between 9 AM and 7 PM and tested 15 min later, whereas subjects in the 9 h group were trained between 9 AM and 11 AM and tested 9 h later. Note that *only the 24 h group underwent a full night of sleep between training and test*.

All volunteers learned to compensate for the optical rotation by the end of training as indicated in Figure 2a. Learning was similar across groups as indicated by the rate of adaptation (F(5,128)=0.85, p=0.52) and the attained level of asymptotic performance during the last block of training (F(5,128)=1.22, p=0.30). As depicted by Figure 2b, memory retention, assessed using error-clamp trials as a percentage of the asymptotic performance, declined progressively over time (F(5,128)=13.75, p<0.001; mean±SEM: 15 min = 79.6±3.1 %; 1 h = 66.8±3.9 %; 3 h = 53.6±4.7 %; 5.5 h = 44.1±4.2 %; 9 h = 42.0±5.6 %; 24 h = 40.5±%4.1 %). This pattern of forgetting conformed to a single exponential function y(t) = a*exp(-b*t)+c (see Supplementary Figure S1), typically observed in declarative and force-field adaptation tasks (Criscimagna-Hemminger & Shadmehr, 2008; Murre & Dros, 2015). Specifically, SMA memory decayed with a time constant of 2.3 h (*b* = 0.43 h^-1^) and reached a plateau at approximately 5.5 h post-learning. The latter was determined based on the time point at which memory retention did not statistically differ from the asymptote *c* obtained from the exponential fit (*c* = 40.67%; retention at 5.5 h vs. *c*, t(21) = 0.807, p = 0.86, whereas retention at 3 h vs. *c*, t(21) = 2.745, p = 0.024). Memory retention stabilized thereafter, with no further decay -nor increment-observed in the 9 h and 24 h groups compared to the 5.5 h group (ANOVA between the 5.5 h, 9 h, and 24 h groups: F(2,146)=0.16, p=0.85).

**Figure 2.**
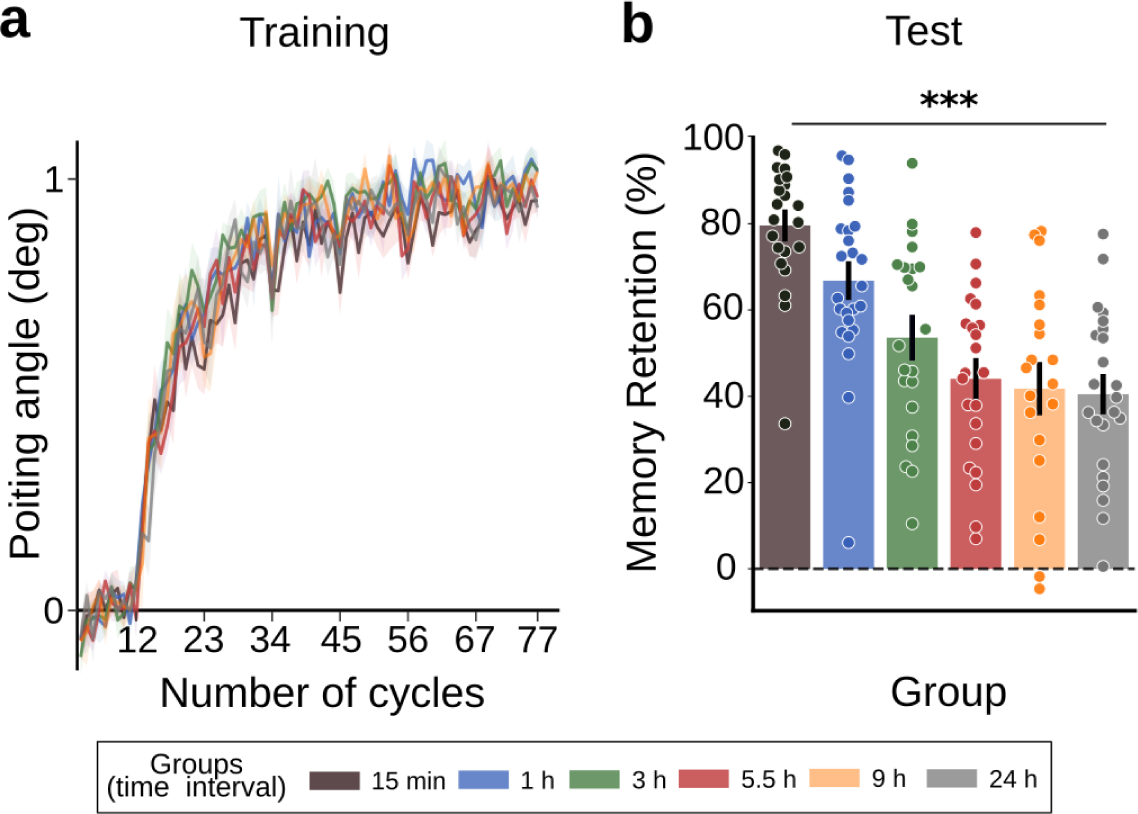
Experiment 1. Sleep does not benefit motor skill maintenance when training occurs far from sleep. Six groups of participants adapted to a CCW 30-degree optical rotation during Training, and memory retention was assessed at different time points during Test, indicated in the legend with colors. During training on the SMA task, all participants performed 1 baseline block of null trials (11 cycles) followed by 6 blocks of perturbed trials (66 cycles). a) Learning curves. Shown are the median±SEM of the pointing angle during visuomotor adaptation for all six groups. b) Memory Retention. Memory retention was evaluated during the test session by quantifying the pointing angle through two error-clamp cycles, and expressed as a percentage of the asymptotic performance level. Shown are the mean±SEM for each group, and the individual data superimposed as dots. *** p<0.001 indicates the result of the one-way ANOVA test across groups.

Altogether, these findings show that motor skill maintenance does not benefit from a full night of sleep when the temporal proximity between training and bedtime is not controlled for. Note that, under these experimental conditions, our results are in line with the prevailing literature supporting the notion that SMA memory consolidates with the passage of time (Donchin et al., 2002; Doyon et al., 2009; Debas et al., 2010; Thürer et al., 2018).

### Motor skill maintenance stabilizes within a ∼6-hour time window

In Experiment 1 we showed that sleep is ineffective when it takes place far from training. We hypothesize that a benefit of sleep would rather be observed when it occurs closely after training, while it overlaps with the memory stabilization window. To determine the optimal time window between learning and sleep, in Experiment 2 we examined the time course of SMA memory stabilization using an anterograde interference protocol (Wigmore et al., 2002; Tong & Flanagan, 2003; Sing & Smith, 2010; Leow et al., 2014). We have previously shown that successive adaptation to opposing optical rotations leads to a deficit in the learning rate when they are separated by 5 min and 1 h. This deficit tends to dissipate around 6 h (Lerner et al., 2020). While our previous study provides evidence on memory encoding, it does not deepen into the process of memory consolidation. To this aim, here we investigated the impact of anterograde interference on previously unpublished long-term memory retention derived from the same dataset.

Four groups of participants (n=15-20 per group) adapted sequentially to two 30-degree opposing optical rotations (A=CCW followed by B=CW) separated by one of four possible time intervals: 5 min, 1 h, 6 h, or 24 h (Figure 1b, middle panel). In addition, a control group (n=20) only adapted to B. Long-term memory for all groups was assessed 24 h after training on B. For practicality, we reproduce the learning curves from our previous work in Figure 3a. Visual inspection suggests that all participants learned to compensate for perturbation A to a similar extent (refer to Lerner et al., 2020, for corresponding statistics regarding the rate of adaptation and achieved level of asymptote). In contrast, anterograde interference significantly affected adaptation to B; while all groups reached asymptotic performance, the 5 min and 1 h groups were significantly slower than the control group (refer to Lerner et al., 2020 for corresponding statistics).

**Figure 3.**
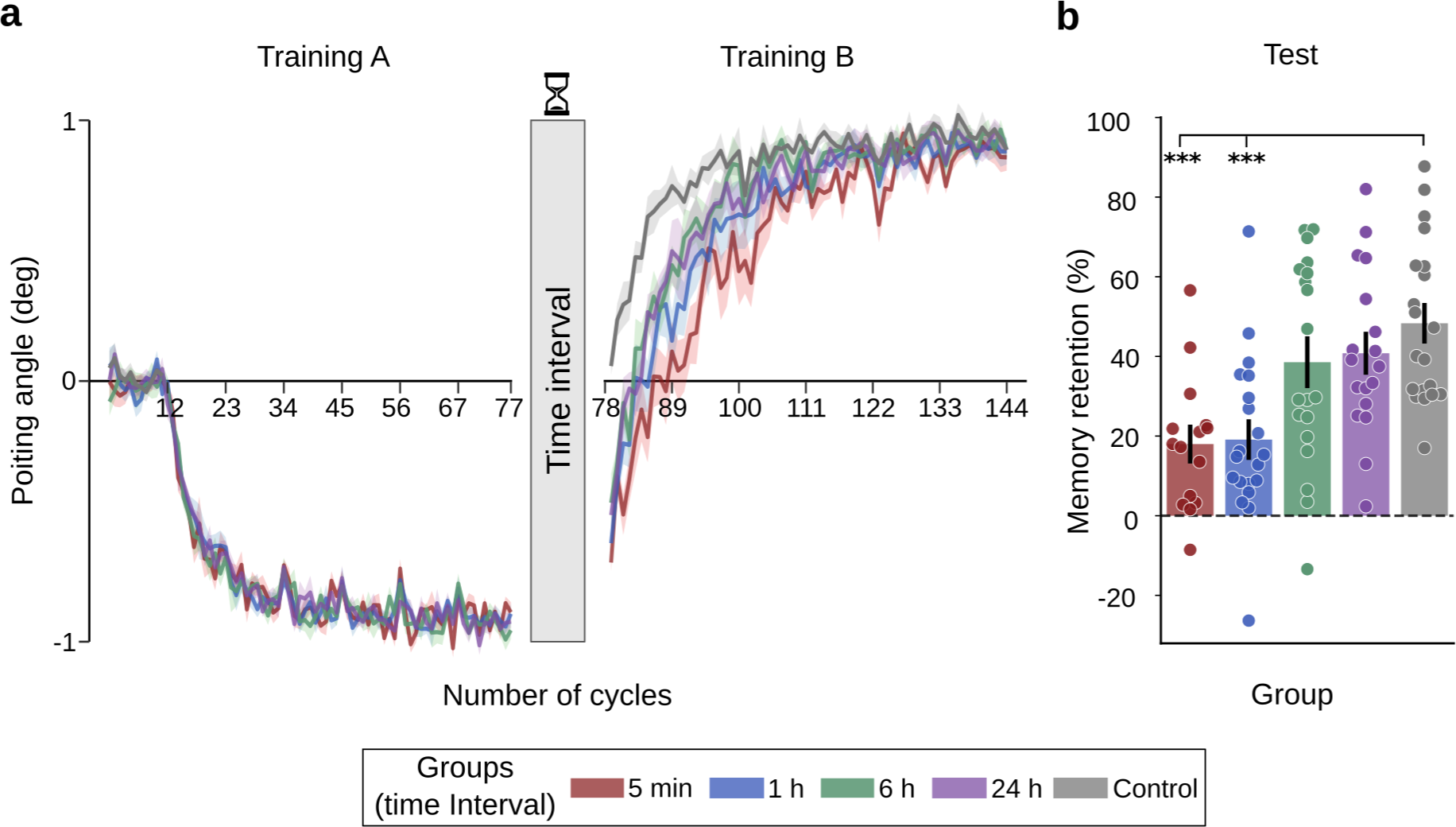
Experiment 2. Motor skill maintenance stabilizes within a ∼6-hour time window. Four groups of participants adapted to two opposing optical rotations (Training A and Training B) separated by different time intervals (indicated in the legend with colors). An additional group (Control) was only trained on B. All 5 groups were tested 24 h after B (Test). During training, all volunteers performed 1 baseline block (11 cycles) of null trials and 6 blocks (66 cycles) of a CCW 30-degree optical rotation (A), followed by 6 blocks of a CW 30-degree rotation (B). a) Learning curves. Shown are the median±SEM of the pointing angle for all groups as a function of the training session. b) Memory retention. Memory retention was evaluated during the Test session by quantifying the pointing angle through two error-clamp cycles, expressed as a percentage of the asymptotic performance on B. Shown are the mean±SEM of memory retention for each group, with individual data superimposed as dots. *** p<0.001 indicates those groups that differed significantly from the control group according to Dunnett’s test. Figure 3a was reproduced from Lerner et al., 2020 with the permission of Oxford University Press.

Regarding the time window of memory stabilization (Figure 3b), here we found that memory retention was significantly hindered by anterograde interference (F(4,87)=7.61, p<0.001; mean±SEM: 5 min = 18.0±4.3 %; 1 h = 19.1±4.5 %; 6 h = 38.6±5.9 %; 24 h = 41.0±4.8; control = 48.3±4.5 %). Specifically, a strong deficit in memory retention was observed at 5 min and 1 h (Dunnett’s test; 5 min vs. control: p<0.001; 1 h vs. control: p<0.001), which dissipated at 6 h (Dunnett’s test; 6 h vs. control: p=0.41; 24 h vs. control: p=0.65). This temporal pattern, which resembles that observed for memory encoding, is consistent with release from interference (Brashers-Krug et al., 1996; Shadmehr & Brashers-Krug, 1997).

One may wonder if the relatively less amount of time spent at the asymptote for the 5 min and 1 h groups, also known as “overlearning” (Krakauer et al., 2005; Shibata et al., 2017; Mooney et al., 2021), may explain the lower level of retention for these groups depicted in Figure 3b. Supplementary Figure S2 rules out this potential confound by demonstrating that an additional group trained on a similar amount of overlearning on B (without previous training on A), achieved a level of memory retention comparable to the control group.

In sum, Experiment 2 reveals that motor skill maintenance memory stabilizes during wake through a ∼6 h time window. This provides experimental evidence supporting memory consolidation of these types of motor memories.

### Sleep benefits motor skill maintenance when it occurs immediately after learning

Building on the findings of Experiment 2, in Experiment 3 we investigated the impact of manipulating the temporal gap between training and sleep on long-term memory and the neurophysiological markers of sleep consolidation. Our working hypothesis posited that sleeping early within the memory stabilization window, while the memory remains fragile, would enhance memory retention through an active mechanism.

To test this hypothesis two groups of volunteers (n=21 and n=23) adapted to a CW 30-degree optical rotation either immediately before or approximately 14 h before a full night of sleep, and memory retention was assessed 24 h later (Figure 1b, bottom panel). Polysomnography recordings were acquired during sleep for these two groups. We refer to the first group, trained and tested at night (∼10 PM), as PM/PM, and to the second group, trained and tested in the morning (∼8 AM), as AM/AM. Two additional control groups were included to eliminate the possibility of a circadian effect at the time of testing: a PM/AM group, trained in the evening but tested in the morning (with an intermediate sleep period), and an AM/PM group, trained in the morning but tested in the evening (without an intermediate sleep period). About ∼20 minutes (mean±SEM = 21.9±2.8 minutes) took place between the end of training and the onset of NREM1 sleep for the two groups that trained PM, confirming a good overlap between sleep and the memory stabilization window.

All four groups adapted similarly to the optical rotation regardless of the time of training (Figure 4a), as determined based on the rate of learning (F(3,70)=1.46, p=0.23) and the achieved asymptotic performance (F(3,70)=0.860, p=0.461). Critical to our manipulation, and in alignment with our hypothesis, we observed a 31% increase in memory retention (Figure 4b) in the groups that trained immediately before sleep (PM/PM and PM/AM) compared to the groups that trained distant from sleep (AM/AM and AM/PM) (two-way ANOVA; significant main effect of training time: F(1,70) = 5.52, p = 0.02; mean±SEM: PM/PM = 55.5±4.4 %; PM/AM = 55.9±10.5 %; AM/AM = 42.9±4.3 %; AM/PM = 41.8±5.6 %). This pattern of memory retention cannot be explained by the time of test (two-way ANOVA; non-significant effect of time of test: F(1,70)=0.18, p=0.67), ruling out a circadian modulation at the level of retrieval.

**Figure 4.**
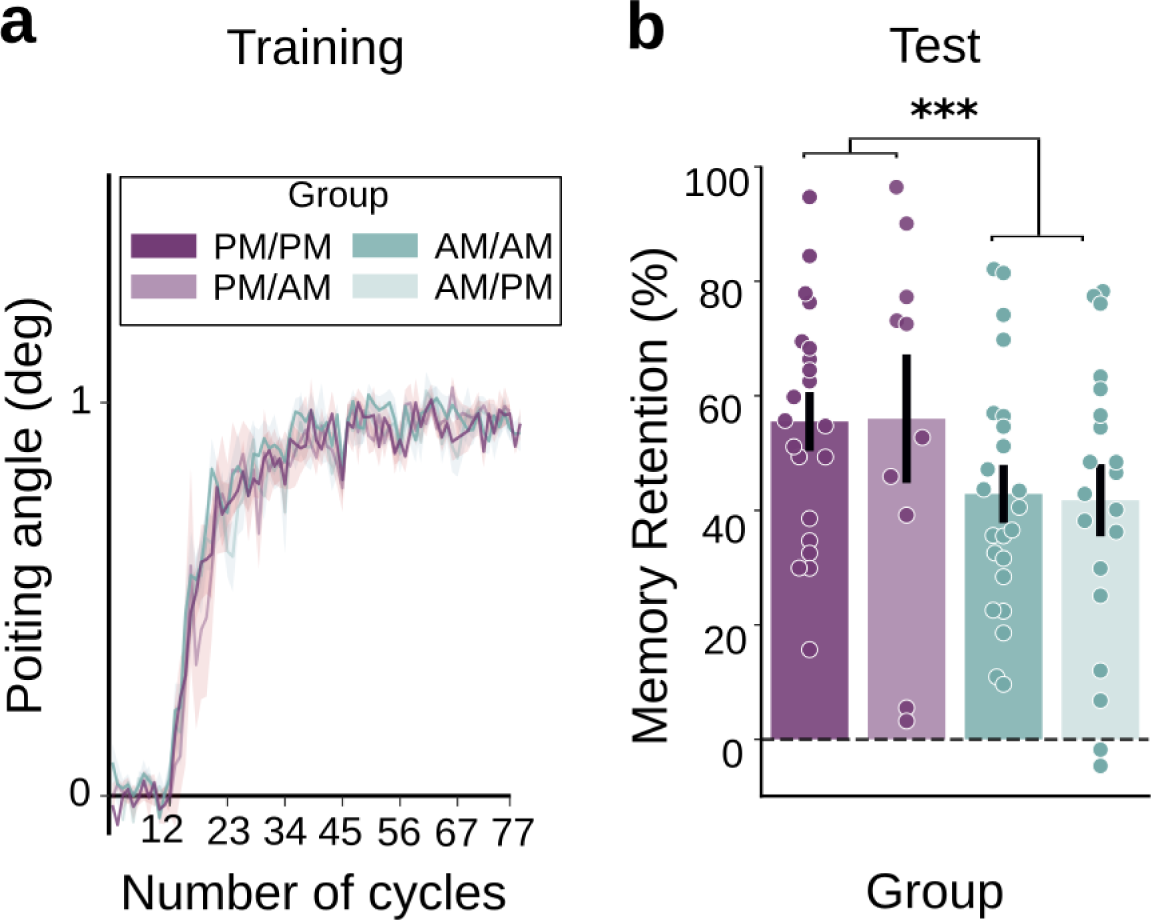
Experiment 3. Sleep potentiates motor skill maintenance when it occurs immediately after learning. Two groups of volunteers trained either far or close to bedtime and were tested 24 h later (AM/AM and PM/PM). To rule out a possible circadian effect due to the time of Test, two additional groups were also trained either far or close to bedtime but were tested 9 h later (AM/PM and PM/AM). During training on the SMA task, all participants performed 1 baseline block of null trials (11 cycles) followed by 6 blocks (66 cycles) of a CW 30-degree optical rotation. a) Learning curves. Shown are the median±SEM of the pointing angle corresponding to the AM/AM and PM/PM groups and their circadian controls (AM/PM and PM/AM). b) Memory retention. Memory retention was evaluated during the Test session by quantifying the pointing angle through two error-clamp cycles, which was expressed as a percentage of the asymptotic performance level. Shown is the mean±SEM of the memory retention attained by each group; individual data is superimposed as dots. *** p<0.001 indicates the result of the two-way ANOVA for the main effect of time of training on memory retention.

Notably, the level of memory retention achieved by the AM/AM group was comparable to that attained by the 24 h group from Experiment 1, which trained sparsely during wakefulness (memory retention, mean±SEM: AM/AM = 42.9±4.3%; 24 h group from Experiment 1 = 40.5±4.1%; t(43.8)=0.40, p=0.3). This result confirms that the beneficial impact of sleep observed herein represents a *net enhancement of long-term memory*.

Our findings indicate that sleeping during the stabilization window potentiates motor skill maintenance. We hypothesized that this effect reflects, at least in part, an active role of sleep in the consolidation of newly acquired information. To test this hypothesis, we assessed the effect of our manipulation on two well-established neural markers of sleep consolidation, which have been consistently observed across species and learning paradigms (Maingret et al., 2016; Ladenbauer et al., 2017; Helfrich et al., 2018; Muehlroth et al., 2019; Navarro-Lobato and Genzel, 2019). We and others have shown that motor learning increases the density of fast spindles and the SO-spindle coupling during NREM sleep when sleep takes place after training (Kim et al., 2019; Silversmith et al., 2020; Solano et al., 2022a; 2022b; Hahn et al., 2022). In our previous studies (Solano et al., 2022a, 2022b), we found that this modulation predominantly occurs over the left hemisphere, contralateral to the trained hand, and predicts overnight long-term memory. Here we analyzed the EEG recordings from the AM/AM and PM/PM groups obtained during the night following SMA learning and contrasted these sleep metrics. We predicted that if sleep actively contributes to the consolidation of newly acquired SMA memory, it should increase the density of sleep spindles and SO-spindle couplings specifically over the contralateral hemisphere *only* in the PM/PM group.

As illustrated in Figure 5a, we found that training close to bedtime increased the spindle density over the left -contralateral-hemisphere (left/right hemisphere % change of fast spindle density, mean±SEM: PM/PM = 12.1±1.6 %, AM/AM = 1.6±1.5 %; F(1,38.87)=6.48, p=0.015, followed by t-test versus zero, PM/PM: t(20.36)=4.1, p=0.001, AM/AM: t(18.8)=0.53, p=1). Likewise, Figure 5b shows that our manipulation also enhanced the spindle-SO coupling over the left hemisphere (left/right hemisphere % change of fast spindle-SO density, mean±SEM: PM/PM = 14.8±1.6 %, AM/AM = 1.6±1.1 %; F(1,37.74)=6.02, p=0.019, followed by t-test versus zero, PM/PM: t(19.8)=3.5, p=0.004, AM/AM: t(19.2)=0.52, p=1). No significant differences were found in sleep architecture across groups suggesting that our results may not be attributed to differences in the quality or duration of sleep (Supplementary Table S1).

**Figure 5.**
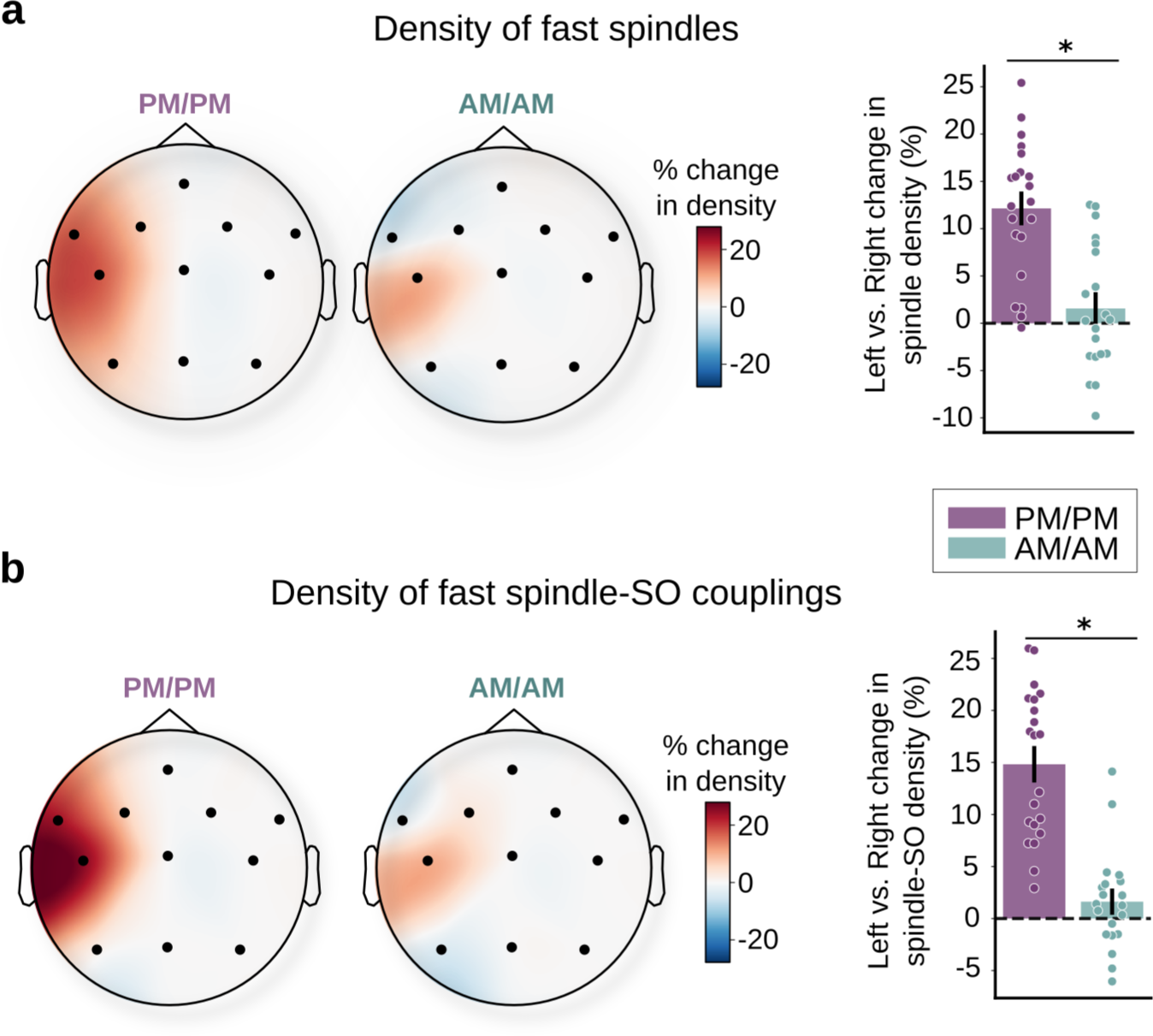
Sleep benefits motor skill maintenance through an active mechanism. Shown are the topographic plots depicting the spatial distribution of the inter-hemispheric percent change of the density of fast spindles (a), and the fast spindle-SO couplings (b) during NREM of the first cycle of sleep for the PM/PM and the AM/AM groups. The inter-hemispheric change in these metrics was computed according to the function (Left Hemisphere-Right Hemisphere)/Right Hemisphere*100, applied across corresponding EEG electrodes. Barplots on the right depict the statistical quantification of these metrics across groups obtained based on LMMs. Superimposed on the bar plots is the estimated grand average for each subject (illustrated as dots). * p<0.05 indicates the result of the F test across groups.

In sum, Experiment 3 indicates that sleep potentiates long-term memory when it occurs shortly after learning, during the window of memory stabilization. The fact that the observed behavioral gain cannot be attributed to circadian effects, along with the specific contralateral modulation in spindle density and spindle-SO coupling only in the group that trained close to bedtime strongly suggests that motor skill maintenance undergoes consolidation during sleep.

## Discussion

While there is compelling evidence that sleep improves different types of memories, its role in motor memory consolidation remains a topic of contention. Current work suggests that motor skill memory requires sleep to consolidate while motor skill maintenance is consolidated with the passage of time, irrespective of sleep. This evidence has led to the proposal that the latter may be an exception to other types of memories (Brodt et al., 2023). In the present study, we addressed this ongoing debate through a series of three meticulously designed experiments. In line with previous work, we show that SMA memory consolidates with the passage of time when training is distributed throughout the daytime. However, when the time interval between learning and bedtime is manipulated to ensure that sleep coincides with the window of memory stabilization, a significant memory enhancement becomes apparent. This marked improvement in long-term memory was accompanied by specific modulation of neural markers of sleep consolidation, including an increase in spindle density and spindle-SO coupling during NREM, thereby providing support for an active role of sleep behind the behavioral benefit.

Our work contributes to reconciling conflicting viewpoints regarding the mechanisms involved in the consolidation of procedural memories, namely MSL vs. SMA. It is important to note that while numerous studies on SMA have provided supporting evidence for the hypothesis that both visuomotor and force-field adaptation memories consolidate over time (Donchin et al., 2002; Doyon et al., 2009; Debas et al., 2010; Thürer et al., 2018), other research conducted by Tononi and colleagues has pointed to a distinct advantage of sleep in the context of visuomotor adaptation (Huber et al. 2004; Landsness et al., 2009). These apparent disparities can be resignified in light of our current results. While in the former set of studies, participants were trained sparsely throughout the daytime, in Tononi’s work volunteers learned the motor task immediately before bedtime. Notably, in none of these studies, the temporal gap between learning and sleep was deliberately considered. By systematically manipulating this gap here we demonstrate a net benefit of sleep on motor memory of approximately 30%. Our findings help settle the above controversy and suggest that, like motor skill learning and declarative learning, motor skill maintenance undergoes sleep consolidation.

Is the sleep-related memory benefit we observe herein the outcome of a passive or an active mechanism? There are currently two main hypotheses supporting an active role of NREM sleep in memory consolidation. According to the systems consolidation hypothesis, newly encoded memories, initially stored in hippocampal networks, are reactivated during slow-wave sleep and gradually integrated with existing memory traces at the level of the neocortex. This process is thought to depend on the close synchrony between slow oscillations, sleep spindles, and hippocampal ripples (Rasch & Born, 2007; Diekelmann & Born, 2010; Buzsáki, 2015; Maingret et al., 2016; Ladenbauer et al., 2017; Latchoumane et al., 2017; Helfrich et al. 2018; Muehlroth et al. 2019; Navarro-Lobato and Genzel, 2019). Another –not mutually exclusive-account of memory consolidation is the synaptic homeostasis hypothesis (SHY), according to which synaptic weights potentiated during learning are downscaled by sleep, improving their signal-to-noise ratio (Tononi and Cirelli 2003, 2006). In line with the systems consolidation hypothesis, here we show that training specifically increased the spindle-SO coupling over the contralateral hemisphere, solely when it occurs immediately before bedtime, supporting an active role of sleep in SMA memory consolidation. Nevertheless, it remains plausible that sleep actively promotes consolidation while also providing passive protection from interference. Although our study was not aimed at testing this possibility, it is a topic of relevance worth future investigation. Note that our experimental design precluded us from directly examining whether our findings align with SHY (but refer to Solano et al., 2022a, for prior evidence from our laboratory supporting both theoretical accounts).

Our findings from Experiment 2 build upon our earlier work, showing that anterograde interference impairs the ability to learn within a 6 h window (Lerner et al., 2020). Here we show further that anterograde interference also hinders long-term memory retention following a similar time course, suggesting that the same biological substrates may support both learning and memory stabilization (Della-Maggiore et al., 2017), a hypothesis in line with the modern concept of an engram (Josselyn & Tonegawa, 2020). Interestingly, SMA memory decay unveiled by Experiment 1, followed a similar temporal evolution to memory stabilization, leveling off around 6 h post-learning, a finding that is in line with work from force-field adaptation (Criscimagna-Hemminger & Shadmehr, 2008). This temporal alignment opens the possibility that the amount of forgetting may depend on the stability of the motor memory trace (Frankland et al., 2013; Davis & Zhong, 2017) so that more stable memories are less likely to undergo forgetting. Further studies in which these behavioral metrics could be tracked in the same participants would be needed to explore this interplay.

Why might the temporal proximity between learning and sleep play a crucial role in the overnight consolidation of SMA memory? While many studies from the declarative and non-declarative memory fields have reported the beneficial effects of aligning learning with sleep (e.g. Gais et al., 2006; Talamini et al., 2008; Doyon et al., 2009; Van Der Werf et al., 2009; Payne et al., 2012; Holz et al., 2012; Inostroza et al., 2013; Sawangjit et al., 2018, 2020; Truong et al., 2023), the precise mechanism/s underlying this phenomenon remain/s elusive. Emerging evidence suggests that the coupling between sleep spindles and slow oscillations (SOs) observed during NREM sleep may promote the occurrence of hippocampal sharp-wave ripples (SWRs), a high-frequency oscillation (∼90Hz) directly implicated in memory reactivation (Ngo et al., 2020; Staresina et al., 2023; Brodt et al., 2023). Notably, this triad (SO-spindle-SWR) has been observed in rodents during the early phases of motor skill learning (Kim et al., 2023). Converging evidence from our lab underscores the involvement of the human hippocampus during the initial phase of MSL, and SMA up to 30 min post-training (Jacobacci et al., 2020; Griffa, G., Jacobacci, F., Della-Maggiore, V., “Unveiling the contribution of the hippocampus to motor learning”, OHBM Meeting 2022). One possibility is that, like declarative learning, the hippocampus enables sleep-dependent motor memory consolidation. However, this process may be initiated only when sleep closely follows learning, ensuring the hippocampus remains actively engaged. Alternatively -but not exclusively-the neurochemical and/or neuromodulatory milieu of sleep may favor the activation of mechanisms associated with memory consolidation (Diekelmann & Born, 2010; Rasch & Born, 2013), such as synaptic homeostasis (Tononi & Cirelli, 2003, 2006, 2014) or *de novo* protein synthesis and gene expression, key for synaptic plasticity (Ramm & Smith, 1990; Nakanishi et al., 1997; Ribeiro et al., 1999; Mackiewicz et al., 2007; Seibt et al., 2012). Rigorous biological interventions would be essential to empirically test these hypotheses that at this stage remain speculative.

In conclusion, our findings indicate that consolidation of motor skill maintenance depends both on the passage of time and sleep. Specifically, we showed that when training is distributed throughout the daytime consolidation proceeds independently of sleep. Conversely, when sleep is strategically scheduled to overlap with the memory stabilization window, SMA memory is enhanced along with a distinct modulation of the neural markers of sleep consolidation. Our work advances research at the basic and translational level. At the basic level, it contributes to resolving a longstanding debate concerning the role of sleep in SMA memory consolidation. Furthermore, it opens the possibility of common mechanisms supporting consolidation across different procedural and declarative memory domains. Finally, at the translational level, it may impact rehabilitation programs, potentially expediting motor injury recovery by aligning training sessions with the sleep cycle or incorporating strategic nap interventions.

## Materials and Methods

### Participants

A total of 270 participants (139 females; mean±SD = 24.3±4 years old) with no known history of neurological or psychiatric disorders were recruited from the School of Medicine of the University of Buenos Aires. Subjects were right-handed as assessed by the Edinburgh Handedness Questionnaire (Oldfield, 1971), and were asked to maintain a regular sleep schedule before and during the study. This was monitored through self-recorded spreadsheets provided by the researcher

All volunteers signed the informed consent approved by the Ethics Committee of the Hospital de Clínicas (University of Buenos Aires), which complies with the Declaration of Helsinki in its latest version, and with the National Law on the Protection of Personal Data.

### Experimental Paradigm

Motor skill maintenance was studied using a visuomotor adaptation SMA task (Figure 1a), which has been previously described in detail elsewhere (e.g. Lerner et al., 2020; Solano et al., 2022a) and is briefly summarized here. Participants performed a center-out task consisting of moving a cursor from a start point in the center of a computer screen to one of eight visual targets arranged concentrically, using a joystick controlled with the thumb and index finger of the right -dominant-hand. The vision of the hand was occluded. Subjects were instructed to perform a shooting movement to one of the eight targets with the cursor, as soon as it appeared on the screen. The order of target presentation was randomized within cycles. One cycle consisted of 8 trials, with a single movement aimed at each target, and each block consisted of 11 cycles.

Three types of trials were presented throughout the study, which varied depending on the experiment (Experiment 1 through 3), the group (experimental or control), and the session (training or test). During null trials, in which no perturbations were applied, the movement of the cursor directly mapped onto the joystick movement. During perturbed trials, a counterclockwise (CCW) or a clockwise (CW) optical rotation of 30 degrees was applied to the cursor, deviating its trajectory. During error-clamp trials (EC), the cursor trajectory was manipulated to provide fake “straight” paths to the target that mimicked those generated during correct trials. The latter was accomplished by projecting the actual movement of the cursor to the straight line between the start point and the target, with some controlled variability (10-degree standard deviation). These trials allowed to measure memory retention without the confound of learning from error (Criscimagna-Hemminger & Shadmehr, 2008). The SMA task was implemented in MATLAB (The MathWorks, Inc.) using the Psychophysics Toolbox v3 (Brainard, 1997).

### Experimental design

#### Experiment 1

The aim of Experiment 1 was to examine the impact of sleep on motor skill maintenance when training occurred far from bedtime. To this aim, we tracked the temporal evolution of memory retention through a 24 h period when training took place throughout the daytime (Figure 1b, upper panel). One hundred and thirty-four (134) participants were randomly assigned to 6 groups defined by the time elapsed between the end of the training session and the test session: 15 min (n = 22), 1 h (n = 25), 3 h (n = 22), 5.5 h (n = 22), 9 h (n = 20), or 24 h (n = 23). Critically, subjects were trained at any time between 9 AM and 7 PM. Note that the 24 h group was the only one that slept between training and test.

During the training session, participants of all groups were initially familiarized with the joystick and the experimental paradigm through one block of null trials (baseline), in which the cursor reliably represented the position of the hand. Then, participants faced six blocks of perturbed trials in which a CCW 30-degree optical rotation was applied to the cursor. During the test session, subjects were exposed to two EC cycles to assess memory retention.

#### Experiment 2

Experiment 2 was designed to establish the time course of memory stabilization during wakefulness, with the ultimate goal of determining the specific time window during which sleep might be most effective in modulating SMA memory (addressed in Experiment 3). To this aim, we analyzed unpublished data acquired as part of a larger study aimed at characterizing the effect of anterograde interference on SMA, some of which we reported recently (Lerner et al., 2020). In our previous work, we showed that adaptation to an optical rotation hinders the ability to adapt to the opposite rotation within a 6 h window. Unlike retrograde interference protocols, which have mostly failed at unveiling the time course of memory consolidation in sensorimotor adaptation (although see Brashers-Krug et al. 1996; Shadmehr & Brashers-Krug, 1997 for exceptions), we showed that the use of an anterograde interference protocol yielded a gradual pattern of release from interference. This suggests that our approach may be a good alternative to track memory consolidation in this type of learning. Thus, here, we analyzed the impact of anterograde interference on long-term memory to directly determine the time window of memory consolidation during wakefulness (Figure 1b, middle panel). After a baseline period consisting of 1 block of null trials, four groups of participants were sequentially exposed to 6 blocks of a 30-degree CCW optical rotation (A) followed at different time intervals by 6 blocks of a 30-degree CW optical rotation (B). The time intervals elapsed between A and B were 5 min (n = 15), 1 h (n = 20), 6 h (n = 19), or 24 h (n = 18). A fifth group, which acted as control (n = 20), trained only on rotation B without being previously exposed to A. All volunteers returned 24 h after adaptation to B for the test session, during which they were exposed to two EC cycles to quantify long-term memory retention. Participants were instructed not to nap between adaptation sessions.

#### Experiment 3

The objective of Experiment 3 was to examine the effect of sleep on motor skill mainenance when the temporal gap between training and bedtime is controlled. We used the results of Experiment 2 to guide the precise temporal proximity between training and sleep. Two different groups of participants were trained on the SMA task many hours prior (∼14 hour) or within an hour before sleep (Figure 1b, bottom panel). We will refer to the former group as AM/AM because volunteers underwent training and testing in the morning, and to the latter as PM/PM because participants in this group were trained and tested at night. Volunteers within the AM/AM group (n=23) came to the laboratory in the morning of Day 1 (∼8-9 AM) to participate in the SMA training session and returned ∼14 h later for a full night of sleep. Conversely, participants within the PM/PM group (n=21) came to the laboratory during the night of Day 1 (∼9-10 PM), participated in the SMA training session, and went to sleep ∼10 min later. Memory retention was assessed during the test session on Day 2, 24 h post-training (∼8-9 AM for the AM/AM group, and around ∼9-10 PM for the PM/PM group). All participants were explicitly instructed to refrain from daytime napping. The SMA training session consisted of one baseline block of null trials followed by six blocks of perturbed trials in which a 30-degree CW optical rotation was imposed on the cursor. During the test session, participants were exposed to 2 cycles of EC trials to assess memory retention. In addition, participants from both the AM/AM and PM/PM groups underwent a polysomnographic (PSG) recording through the full night of sleep (see detailed description below). Only subjects fulfilling the criteria for good sleep quality based on the Pittsburgh Sleep Quality Questionnaire (Buysse et al., 1989) and the Epworth Drowsiness Scale (Johns, 1991) were included in the experiment.

To control for a potential circadian modulation associated with the time of test, two additional groups of participants were included in the statistical analysis. An AM/PM group (n = 20), was trained in the morning and tested the same night (∼9 h later) without intermediate sleep, whereas a a PM/AM group (n = 10) was trained at night and tested the next morning (∼9 h later), after a night of sleep. Data from these groups was reported in our previous work (Solano et al., 2022a).

### PSG Recording

Eleven surface EEG electrodes were placed over the prefrontal, motor, and parietal areas (FC1, FC2, FC5, FC6, C3, C4, P3, and P4) and over the midline (Fz, Cz, and Pz). Electrodes were mounted following the standard 10-20 arrangement (Modified Combinatorial Nomenclature; Oostenveld & Praamstra, 2001). Both mastoids were used as references. In addition to EEG electrodes, two electrodes were placed over the periorbital area of both eyes and two additional electrodes over the chin to measure electrooculography (EOG) and electromyography (EMG), respectively. All signals were acquired at 200 Hz, using the Alice 5 (Philips Respironics, PA, EEUU) or BWmini (Neurovirtual, FL, EEUU) devices.

### EEG Processing

EEG, EOG, and EMG signals were bandpass-filtered to facilitate sleep scoring (EEG: 0.5–30 Hz; EOG: 0.5–15 Hz; EMG: 20–99 Hz). All PSG recordings were sleep-staged manually, according to standard criteria (Iber, 2004). Namely, 30-second epochs were classified as either Wake (W), Non-Rapid Eye Movement (NREM1, NREM2, and NREM3), or Rapid Eye Movement (REM) stage. After stage classification, sleep architecture was determined based on the following measures, expressed in minutes: total sleep time, sleep latency (latency to NREM1), REM latency, total wake time, wake after sleep onset (WASO), and time in NREM1, NREM2, NREM3, and in REM. Sleep efficiency was also computed as the percentage of total sleep time relative to the time interval between lights-off and lights-on (%). Movement artifacts on the filtered EEG signal were detected by visual inspection and manually rejected.

Slow Oscillations (SOs, 0.5–1.25 Hz) and sleep spindles (10–16 Hz) were automatically identified from the EEG signal corresponding to the stages NREM2 and NREM3 by using previously reported algorithms (see below).

#### Detection of SOs

The algorithm implemented to detect SOs was based on that reported by Mölle and collaborators (2011) and Antony and Paller (2017), and it is the same that we used in previous works (Solano et al., 2022a, 2022b). The EEG signal was bandpass-filtered between 0.5 and 1.25 Hz. To quantify SOs, we first identified zero crossings of the EEG signal and labeled them as positive-to-negative (PN) or negative-to-positive (NP). Those EEG segments between two NP zero crossings were considered SOs if they lasted between 0.8 and 2 seconds. Next, we computed the peak-to-peak (P-P) amplitude as the difference between the positive peak and the negative peak. Finally, we determined the median of the P-P amplitudes for each channel and each subject and retained those SOs with a P-P amplitude greater than the median value (Mizrahi-Kliger et al., 2018).

#### Sleep Spindles Detection

The algorithm implemented to detect sleep spindles was based on that reported by Ferrarelli et al. (2007) and Mölle and collaborators (2011), and it is the same that we used in previous works (Solano et al., 2022a, 2022b). The algorithm was run for each channel and each subject. First, the EEG signal was bandpass-filtered between 10 and 16 Hz before calculating the instantaneous amplitude (IA) and instantaneous frequency by applying the Hilbert Transform (Tort et al., 2010). The IA was used as a new time series and was smoothed with a 350 milliseconds moving average window. Next, those segments of the IA signal that exceeded an upper magnitude threshold (90th percentile of all IA points) were labeled as potential spindles. The beginning and end of potential spindles were defined as the time points at which the signal dropped below a lower threshold (70th percentile of all IA points). Putative spindles with a duration between 0.5 and 3 seconds were labeled as true spindles. Finally, only fast spindles with a mean frequency ≥12 Hz (Mölle et al. 2011; Cox et al. 2017) were included in further analysis, as they have been linked to memory consolidation during sleep (Barakat et al., 2011; Ladenbauer et al., 2017, Helfrich et al., 2018; Muehlroth et al., 2019; Navarro-Lobato and Genzel, 2019; Solano et al., 2022a, 2022b).

#### Coupling between SOs and Spindles

After identifying spindles and SOs, we looked for spindles that occurred during a SO. We quantified spindle–SO couplings according to the following criterion: if a spindle had its maximum P-P amplitude within ±1.2 second around the trough of a SO, it was counted as a spindle–SO coupling (Muehlroth et al. 2019; Kurz et al., 2021; Solano et al., 2022a). This algorithm was applied to each channel of each session.

### Data Analysis

#### Behavior

Behavioral performance was assessed based on the pointing angle, which was computed for each trial as the angle of motion of the joystick relative to the line segment connecting the start point and target position (e.g., Lerner et al., 2020; Solano et al., 2022a). Trials in which the pointing angle exceeded 120° or deviated by more than 45° from each cycle’s median, were excluded from further analysis. Trial-by-trial data was next converted to cycle-by-cycle time series by computing the median pointing angle across the 8 trials of a cycle, for each individual subject. For the purpose of graphical representation, the pointing angle was normalized when required. We empirically quantified each subject’s learning rate by fitting a single exponential function (y(t) = a*exp(−b*t) + c) to the sequence of pointing angles, where y(t) represents the pointing angle on cycle *t*, *a* and *c* the initial bias and the asymptote of the exponential respectively, and parameter *b* represents the learning rate.

To assess memory retention, the median pointing angle corresponding to each EC cycle was computed and expressed for each subject as a percentage of the asymptotic pointing angle, calculated based on the median of the last block of learning. Finally, the percentage measure was averaged across EC cycles.

#### Memory decay

To characterize the memory decay of motor skill maintenance as a function of time, we fitted a single exponential function (y(t) = a*exp(-b*t) + c) to the memory retention values across individual subjects from the six groups of Experiment 1. Here, y(t) represents memory retention at minute *t*. Parameter *a* represents the initial retention value, *b is* the rate of memory decay and *c* is the asymptote of the function. Exponential functions have been previously used to characterize forgetting in force-field adaptation (Criscimagna-Hemminger & Shadmehr, 2008) and also in declarative tasks (e.g. Wixted, 2004).

#### Electroencephalographic Signal

As described above, SOs and spindles were automatically identified from the EEG signal previously classified as NREM2 and NREM3, corresponding to the first cycle of sleep. We computed the density of fast spindles during NREM sleep (number of fast spindles per minute of NREM sleep) and the density of fast spindles coupled with an SO (number of spindle-SO couplings per minute of NREM sleep).

In our previous work (Solano et al., 2022a), we found that visuomotor adaptation increased the overall density of fast spindles and fast spindles coupled with an SO rather locally, over the hemisphere contralateral to the trained hand (left hemisphere in our experiment). In that work, all volunteers underwent two PSG recordings, one after a control session in which subjects performed the SMA task in the absence of the optical rotation, and another after a SMA learning session in which an optical rotation was applied. The strong inter-hemispheric modulation of the sleep metrics mentioned above was revealed after computing the percent change of these metrics in the experimental session relative to the control session. In the present study, however, participants in the PM/PM and AM/AM groups underwent a unique PSG recording after a SMA learning session. Thus, to assess the effect of learning on the density of sleep spindles and the spindle-SO coupling for the PM/PM and AM/AM groups we computed their inter-hemispheric percent change according to the function (Left Hemisphere-Right Hemisphere)/Right Hemisphere*100. This function was applied for each subject across corresponding EEG electrodes (FC1-FC2, FC5-FC6, C3-C4, and P3-P4). To illustrate the spatial distribution of the percentage difference between hemispheres we report the results in topographic maps (MNE-Python; Gramfort et al. 2013).

### Statistical analysis

Parametric statistics were used to analyze all metrics of interest. Analyses were carried out using R (version 3.6.3; R Core Team, 2017) in RStudio (Rstudio Team, 2015). Statistical differences were assessed at the 95% level of confidence (alpha = 0.05).

For between-subjects statistical comparisons, we used one-way or two-way ANOVA. The variables of interest were either memory retention, the rate of learning (*b* parameter of the exponential function fitted to pointing angle), or the median pointing angle from the last block of adaptation. The fixed factors were the group and the condition associated with the proximity between learning and sleep.

For the sleep metrics, we fitted a Linear Mixed Model (LMM) in which random intercepts were estimated for each subject to take into account the repeated measures. The variable of interest was the inter-hemispheric percent change of the sleep metrics computed for all corresponding EEG electrodes (FC1-FC2, FC5-FC6, C3-C4, and P3-P4), and the fixed factor was the group. To assess the statistical significance of the fixed factor we used F tests with Kenward-Roger’s approximation of the degrees of freedom to obtain p-values (Halekoh & Højsgaard, 2014).

We used the Dunnett’s test or t-tests corrected for multiple comparisons using Bonferroni, for post-hoc assessment.

## Acknowledgments

We thank the Argentinian Ministry of Defense (PIDDEF-2014-2017/#17), and the Argentinian Agency for the Promotion of Science and Technology (FONCyT: PICT2015-0844; PICT2018-1150) for their financial support.

