## Supplementary Information for "Sleep consolidation potentiates skill maintenance"

### Supporting Information

#### Supplementary Figure S1

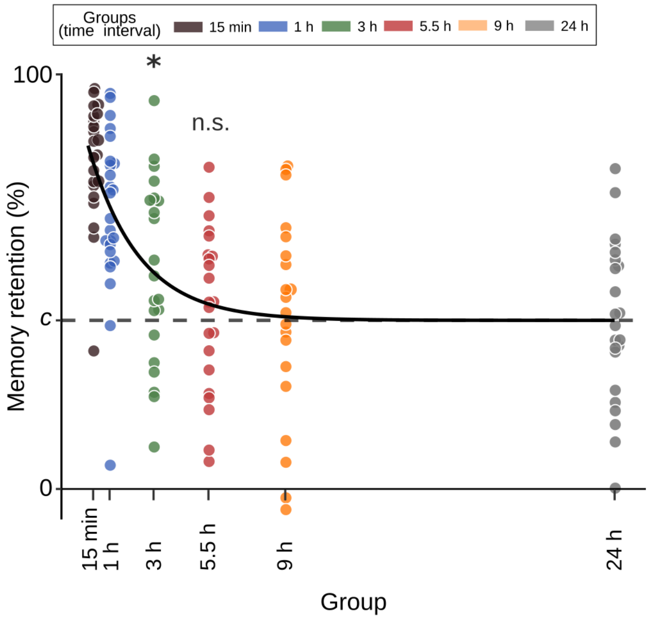


**Supplementary Figure S1. Motor skill maintenance memory decays exponentially.** Shown is SMA memory retention from individual subjects corresponding to each group of Experiment 1. Memory retention was evaluated during the test session by quantifying the pointing angle through error-clamp cycles, which was expressed as a percentage of the asymptotic performance level. The abscissa scale represents the time interval elapsed between the end of training and test (15 min through 24 h). Superimposed is the curve resulting from fitting a single exponential function: y(t) = a*exp(-b*t)+c to memory retention across groups, with *a*=42.47 %, *b*=0.43 $h^{-1}$, and *c*=40.67 %. The dashed line represents the asymptote *c*. Memory decay stabilized ~5.5 h after training. * p<0.05, n.s.: non-significance, indicates the result of the t-test between the 3 h and 5.5 h groups vs. *c* adjusted for multiple comparisons based on Bonferroni.

**Supplementary Figure S2**

In Experiment 2, we showed a significant reduction in memory retention when successive adaptation to optical rotations A and B were separated by 5 min or 1 h. Even though all participants reached the same level of asymptote by the end of training on B, these two groups exhibited a slower rate of learning and, thus, spent less time training at the asymptote than the others. To explore the possibility that this relatively less amount of "overlearning" (Shibata et al., 2017) may explain the pattern of memory retention observed in Figure 3b, we tested an additional group of subjects that trained a similar amount of time at the asymptote as the 5 min and 1 h groups. For practicality, we refer to this group as the overlearning group. We hypothesized that if the observed decrease in memory retention was due to a lesser amount of overlearning rather than an impairment of the memory consolidation process, then the overlearning group should exhibit a similar level of retention to the 5 min and 1 h groups. To determine the amount of training for this additional group we first estimated the overlearning on B as the number of cycles spent training after reaching 95% of asymptotic performance. The latter was estimated based on $3*\tau$, where $\tau = 1/b$ (tau in cycles) is the time constant derived from the single exponential fit applied to the pointing angle and *b* is the learning rate. Thus, the number of cycles training at the asymptote was computed as the total amount of cycles minus $3*\tau$. This yielded the following amount of overlearning: (mean±SEM) 5 min group = 28.2±4.2 cycles; 1 h group = 27.4±4.5 cycles; 6 h group = 36±4.1 cycles; 24 h group = 31.3±4.5 cycles; control group = 43.33±3.3. Given that the 5 min and 1 h groups underwent approximately 2 blocks (around 17 cycles) less training at the asymptote than the control group, the overlearning group (n = 20) trained only on task B for 4 blocks (44 cycles), and was tested for memory retention 24 hours later. As expected, the actual amount of time spent training at the asymptote for the overlearning group (mean±SEM = 24.1±2.4) was similar to the 5 min and 1 h groups ($F(2,52)=0.326$, $p=0.72$).

Supplementary Figure 2 contrasts the amount of overlearning (a), and the level of memory retention attained at 24 h (b) for all experimental groups against the control. Note that, although the overlearning group spent less time training at the asymptote than the control group ($F(3,71)=5.78$, $p=0.001$; followed by Dunnett's test: 5 min vs. control, $p = 0.02$; 1 h vs. control, $p=0.006$; overlearning group vs. control, $p<0.001$), it attained a similar level of retention ($F(3,71)=20.04$, $p<0.001$; followed by Dunnett's test: 5 min vs. control, $p<0.001$; 1 h vs. control, $p<0.001$; overlearning vs. control, $p=0.44$). Collectively, these results confirm that Figure 3b reflects the temporal pattern of SMA memory consolidation.

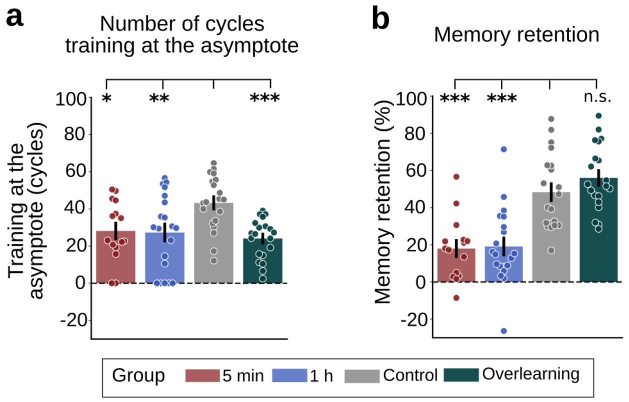


**Supplementary Figure S2. Time course of SMA memory retention is not explained by the time spent training at the asymptote.** Shown are the mean±SEM of the number of cycles training at the asymptote (a), and the level of memory retention (b) for the 5 min, 1 h, and control group from Experiment 2, and the overlearning group. * p<0.05, ** p<0.01, *** p<0.001, n.s.: non-significance, indicate the result of the Dunnett's test for each group compared against the control group.

**Supplementary Table S1. Sleep architecture for AM/AM and PM/PM groups from Experiment 3**

| | AM/AM | | PM/PM | | t-test | |
|---|---|---|---|---|---|---|
| Measure | Mean | SD | Mean | SD | t | p |
| Total Sleep Time (min) | 433.07 | 59.61 | 434.59 | 36.09 | 0.10 | 0.92 |
| Sleep Efficiency (%) | 90.32 | 6.59 | 90.93 | 6.17 | 0.31 | 0.76 |
| Sleep latency (min) | 18.12 | 8.82 | 26.02 | 24.42 | 1.39 | 0.17 |
| REM latency (min) | 117.52 | 45.84 | 99.54 | 36.38 | -1.41 | 0.17 |
| Total Wake Time (min) | 47.74 | 34.32 | 43.23 | 29.24 | -0.46 | 0.65 |
| Wake After Sleep Onset (min) | 20.71 | 24.58 | 15.59 | 15.95 | -0.80 | 0.43 |
| NREM1 (min) | 50.02 | 25.36 | 36.64 | 12.48 | -1.76 | 0.09 |
| NREM2 (min) | 152.19 | 38.31 | 160.88 | 43.02 | 0.69 | 0.49 |
| NREM3 (min) | 117.95 | 33.07 | 127.23 | 30.12 | 0.95 | 0.35 |
| REM (min) | 112.90 | 39.85 | 109.83 | 28.16 | -0.29 | 0.77 |

Shown are the mean and SD corresponding to the sleep measures listed in the first column corresponding to the AM/AM and PM/PM groups from Experiment 3. The last columns depict the statistic and p values yielded from comparing the two groups with t-tests. All measures are depicted in minutes except for sleep efficiency, defined as the percentage of total sleep time relative to the time interval between lights-off and lights-on (%). As observed, no differences were observed in sleep architecture across groups.
